# Enhanced reinstatement of naturalistic event memories due to hippocampal-network-targeted stimulation

**DOI:** 10.1101/2020.08.18.256008

**Authors:** Melissa Hebscher, James E. Kragel, Thorsten Kahnt, Joel L. Voss

## Abstract

Episodic memory involves the reinstatement of distributed patterns of brain activity present when events were initially experienced. The hippocampus is thought to coordinate reinstatement via its interactions with a network of brain regions, but this hypothesis has not been causally tested in humans. The current study directly tested the involvement of the hippocampal network in reinstatement using network-targeted noninvasive stimulation. We measured reinstatement of multi-voxel patterns of fMRI activity during encoding and retrieval of naturalistic video clips depicting everyday activities. Reinstatement of video-specific activity patterns was robust in posterior-parietal and occipital areas previously implicated in event reinstatement. Theta-burst stimulation targeting the hippocampal network increased video-specific reinstatement of fMRI activity patterns in occipital cortex and improved memory accuracy relative to stimulation of a control out-of-network location. Furthermore, stimulation targeting the hippocampal network influenced the trial-by-trial relationship between hippocampal activity during encoding and later reinstatement in occipital cortex. These findings implicate the hippocampal network in the reinstatement of spatially distributed patterns of event-specific activity, and identify a role for the hippocampus in encoding complex naturalistic events that later undergo cortical reinstatement.

## Introduction

Episodic memory allows us to mentally travel back in time to re-experience past events in detail. A prevailing view is that such mental time travel involves reinstating patterns of neural activity that were present during the encoding of an event in a process mediated by the hippocampus [1–3]. The hippocampus is thought to bind together episodic information distributed throughout the brain during encoding and to coordinate reinstatement of this distributed activity at retrieval [4–6]. The current study evaluated this hypothesis by testing whether modulating the hippocampal network via noninvasive brain stimulation affects the strength of cortical reinstatement, measured using functional magnetic resonance imaging (fMRI).

Episodic memory is related to reinstatement of cortical activity as measured by fMRI in humans. These experiments have used multivariate measures [7] to quantify the similarity of multi-voxel activity patterns between initial encoding and subsequent retrieval [8]. This metric of memory-related activity reinstatement is typically robust in occipitotemporal and parietal cortex [4–6,9–11], within areas thought to be involved in representing the sensory features comprising the reactivated memory [3]. The relevance of this metric of reinstatement to real-world memory function has recently been demonstrated by findings of event-specific reactivation of complex video clips that approximate the content of naturally occurring events [9–11]. Some experiments have further found that trial-by-trial fluctuations in hippocampal activity are related to corresponding fluctuations in cortical reactivation strength [4–6], supporting the idea that the hippocampus is important for coordinating distributed reinstatement during episodic memory.

The hippocampus accomplishes its memory-related functions via interactions with distributed networks of interconnected brain regions [5,12–16]. A number of studies have identified a core network of functionally connected regions important for episodic memory including hippocampus, parahippocampal cortex, medial and lateral parietal cortex, and medial prefrontal cortex [13,17–19], although the specific regions that interact with the hippocampus can vary with task and representational demands [20,21]. This core hippocampal network can be functionally influenced using a transcranial magnetic stimulation (TMS) approach referred to as hippocampal-network-targeted (HNT) stimulation. In this approach, TMS is applied to subject-specific cortical locations of the core network defined based on their high functional connectivity to the hippocampus [22]. HNT stimulation using a theta-burst pattern has been shown to enhance performance on classically hippocampal-dependent word-list [23] and spatial precision [24] tests of episodic memory, with associated changes in resting-state fMRI connectivity between the hippocampus and regions of its core network [22,23]. Notably, although previous studies have demonstrated that HNT stimulation can alter hippocampal-network function and episodic memory [22,25,26], the neural mechanisms by which this alteration occurs are unclear. In particular, the impact of HNT on cortical reinstatement of event-specific activity patterns during retrieval has not been tested.

The current study used HNT stimulation to test theorized hippocampal contributions to cortical reinstatement of naturalistic event memories. Participants received theta-burst stimulation to a lateral parietal region defined by high functional connectivity with the hippocampus (HNT stimulation) and then immediately completed a naturalistic video memory task during fMRI scanning. On a different day, the same procedure was performed but with stimulation of a control out-of-network location (vertex). On each day, memory encoding occurred immediately after stimulation during fMRI scanning. Participants viewed a series of short video clips depicting everyday activities, which afford high experimental control while more closely approximating memory for life events compared to traditional laboratory memory tasks. At retrieval, subjects used recall cues to mentally replay studied video clips. To enable comparison with previous episodic and autobiographical memory studies, we measured both objective and subjective memory for video clips (See Figure 1 for experimental design).

**Figure 1.**
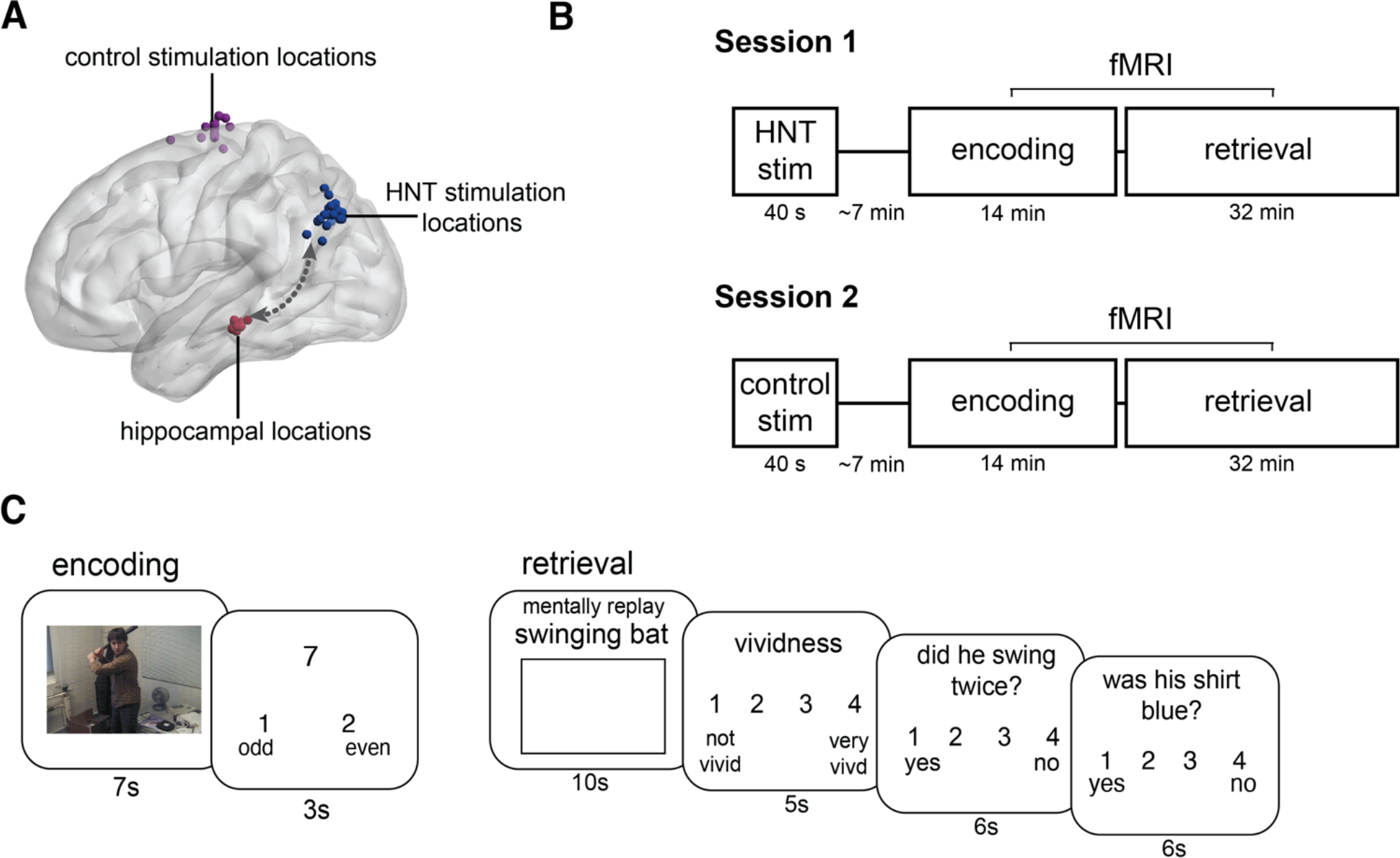
Experimental design overview. (A) HNT stimulation was delivered to subject-specific locations in lateral parietal cortex (blue spheres) defined based on resting-state fMRI connectivity (arrow) with anatomically defined subject-specific hippocampal locations (red spheres). Subject-specific vertex control sites (purple spheres) were anatomically defined in the interhemispheric fissure. (B) In each of two distinct experimental sessions, participants first received stimulation (HNT or vertex control in counterbalanced order across participants). Immediately following stimulation, participants performed a memory task during fMRI. (C) Participants watched a series of videos (51 per session) at encoding and made odd/even judgments to numbers following each video to limit continued processing of videos during the intertrial interval. At retrieval, participants were cued with titles describing each of the studied videos. Subjects mentally replayed each video while viewing a blank screen. They then rated the vividness of the memory and answered two true/false (yes/no) questions to test memory accuracy, including a 4-point confidence judgment.

We measured reinstatement of individual video clips by comparing patterns of fMRI activity between encoding and retrieval [7]. Importantly, the video clips used here were unique from those used in previous studies to identify cortical reinstatement [9,27] in that the clips had relatively high feature overlap, including the same actor, similar contexts, and similar content. This is similar to natural memory demands in which specific events must be remembered given the consistent actors, locations, and objects typical of one’s everyday life. Further, relatively high trial counts (51 trials per session) supported event-specific analyses that allowed us to examine how similarities between neural activity associated with different videos at encoding influences later reinstatement of distinct, event-specific patterns of neural activity.

Based on prior findings of memory enhancement by HNT theta-burst stimulation [23,24] we predicted that, compared to control stimulation, HNT stimulation would improve memory accuracy and increase reinstatement of stimulus-specific activity patterns. Based on evidence for the impact of HNT stimulation on hippocampus [22] and the putative role of hippocampus in coordinating cortical reinstatement [4–6], we hypothesized that HNT stimulation would affect hippocampal activity at encoding and thus influence cortical reinstatement during subsequent retrieval.

## Results

### Effects of stimulation on memory

We first examined the effects of HNT stimulation on memory for videos. Previous experiments have demonstrated improved memory accuracy due to HNT stimulation [22], and we tested for similar effects here by asking two objective yes/no questions about each video clip (Figure 1C). To assess the influence of stimulation on these two questions, we conducted a 2×2 repeated-measures ANOVA with accuracy as the dependent variable, and question number (Q1 vs Q2) and stimulation (HNT vs control) as factors (Figure 2A). There was a significant main effect of question number (F(1,57) = 4.81, p = .033, η^2^_p_ = .078), no main effect of stimulation on accuracy (F(1,57) = 1.15, p = .288, η^2^_p_ = .020), and a significant stimulation by question number interaction (F(1,57)=6.09, p=.017, η^2^_p_ = .097). Paired-samples t-tests were used to determine the effects that led to this interaction. Compared to Q1, Q2 accuracy was significantly worse for control stimulation (t(19) = −3.40, p = 0.003, *d* = 0.838) but not for HNT stimulation (t(19) = 0.21, p = .833, *d* = 0.045). This indicates that the typical pattern of performance on the task (under control stimulation) was a performance decline from Q1 to Q2, and that HNT stimulation rescued this decline. Indeed, HNT stimulation significantly improved Q2 accuracy relative to control stimulation (t(19) = 3.20, p = 0.005, *d* = 0.751), but had no effect on Q1 accuracy relative to control stimulation (t(19) = −0.74, p = 0.470, *d* = −0.206). All paired-sample t-tests reported as significant passed Benjamini-Hochberg correction for multiple comparisons. Difference scores calculated for each participant demonstrated that 15 out of 20 subjects had higher Q2 accuracy for HNT versus control stimulation, and only 3 out of 20 subjects showed a moderate decrease in Q2 accuracy for HNT versus control stimulation, suggesting that this effect is relatively consistent across subjects (Figure 2B). Overall, these findings suggest that HNT stimulation improved the persistence of the memory after retrieval relative to a typical pattern of decline after retrieval evident under control stimulation.

**Figure 2.**
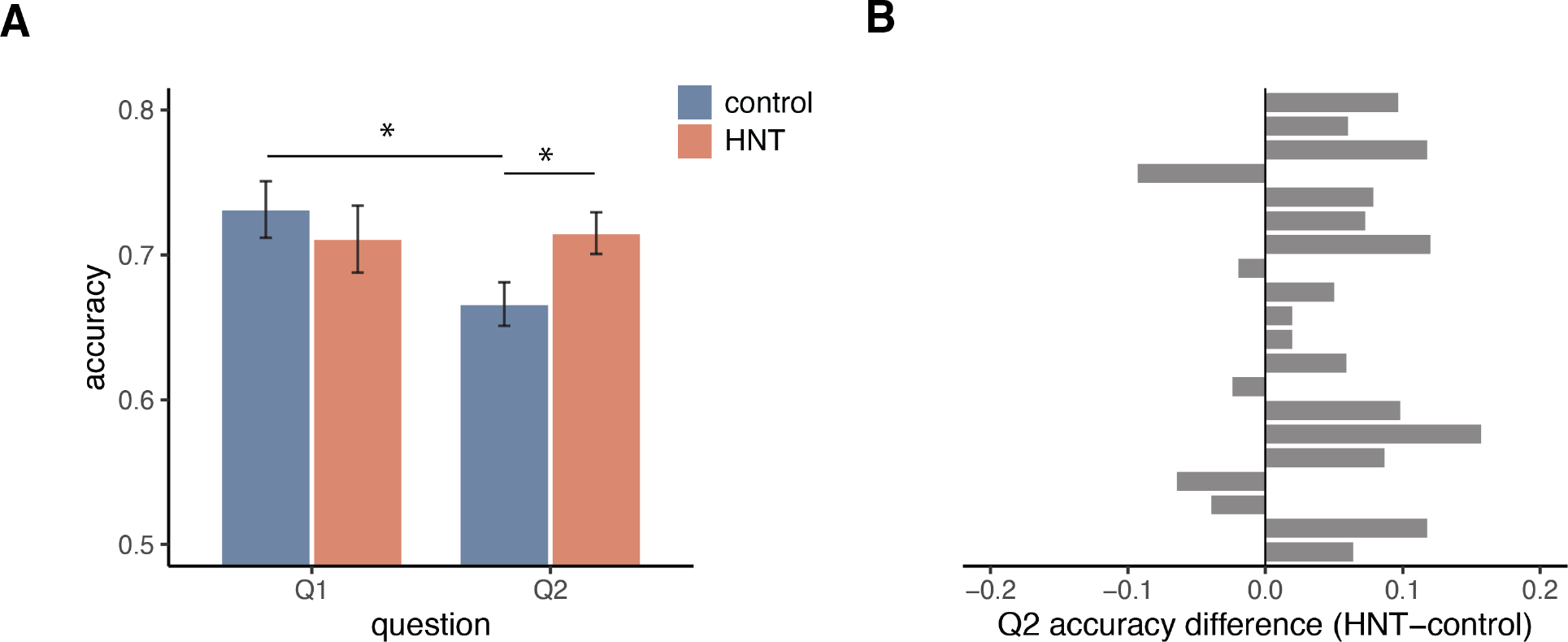
Effects of HNT stimulation on video memory accuracy. (A) Accuracy by question number and stimulation session (HNT and control stimulation). Accuracy on Q2 was significantly better for HNT (red) than control (blue) stimulation. For control stimulation, accuracy was significantly worse on Q2 than on Q1, but there was no difference between questions for HNT stimulation sessions. Error bars represent standard error of the mean. Asterisks indicate tests that were significant after correction for multiple comparisons (p < .05). (B) Difference scores for Q2 accuracy (HNT – control) by participant, showing a relatively consistent improvement in Q2 accuracy for HNT versus control stimulation.

To assess subjective qualities of memory, we measured vividness and confidence (See Episodic Memory Task in Methods). There was no effect of HNT relative to control stimulation on vividness or on confidence for either question (all p’s > .49; See Table 1). Consistent with previous findings and interpretations[22], this suggests that HNT stimulation affects hippocampal-related function, as indicated by improved memory accuracy, more than parietal-related functions, as indicated by no change in memory vividness and confidence judgments.

**Table 1.** Objective and subjective memory assessment values.

| Measure | Mean (SD) |  | t-value |
| --- | --- | --- | --- |
|  | Control | HNT |  |
| Accuracy Q1 | 0.73 (.09) | 0.71 (.10) | -0.76 |
| Accuracy Q2 | 0.67 (.07) | 0.72 (.06) | 3.24* |
| Vividness | 2.71 (.36) | 2.68 (.37) | -0.49 |
| Confidence Q1 | 1.60 (.19) | 1.63 (.17) | 0.70 |
| Confidence Q2 | 1.16 (.23) | 1.16 (.24) | 0.01 |

*t-value reflects HNT vs. control stimulation*
\* $p < .01$

### Effects of stimulation on reinstatement

We next tested the effect of HNT stimulation on reinstatement, which we defined as the video-specific similarity between encoding and retrieval fMRI activity patterns [4,7,9] within cortical regions of interest (ROIs). ROIs were chosen based on their identification in a previous video reinstatement study [9], and included precuneus, angular gyrus, middle occipital gyrus, middle temporal gyrus, and fusiform gyrus in each hemisphere (Figure 3D). For each ROI, encoding-retrieval similarity was computed as the Pearson correlation of a video’s fMRI activity pattern during encoding with the corresponding activity pattern when subjects attempted to mentally replay the same video during retrieval. Encoding-retrieval similarity was computed for all pairs of videos and the average Fisher-transformed correlation of all mismatched pairs (different videos at encoding versus retrieval) was subtracted from the mean correlation of all matched pairs (same video at encoding and retrieval), and z-scored against a null distribution of correlation difference scores obtained by permuting video labels (See Encoding-retrieval reinstatement in Methods). The resulting z-scores reflect pattern similarity, or reinstatement, of video-specific activity for a particular ROI.

**Figure 3.**
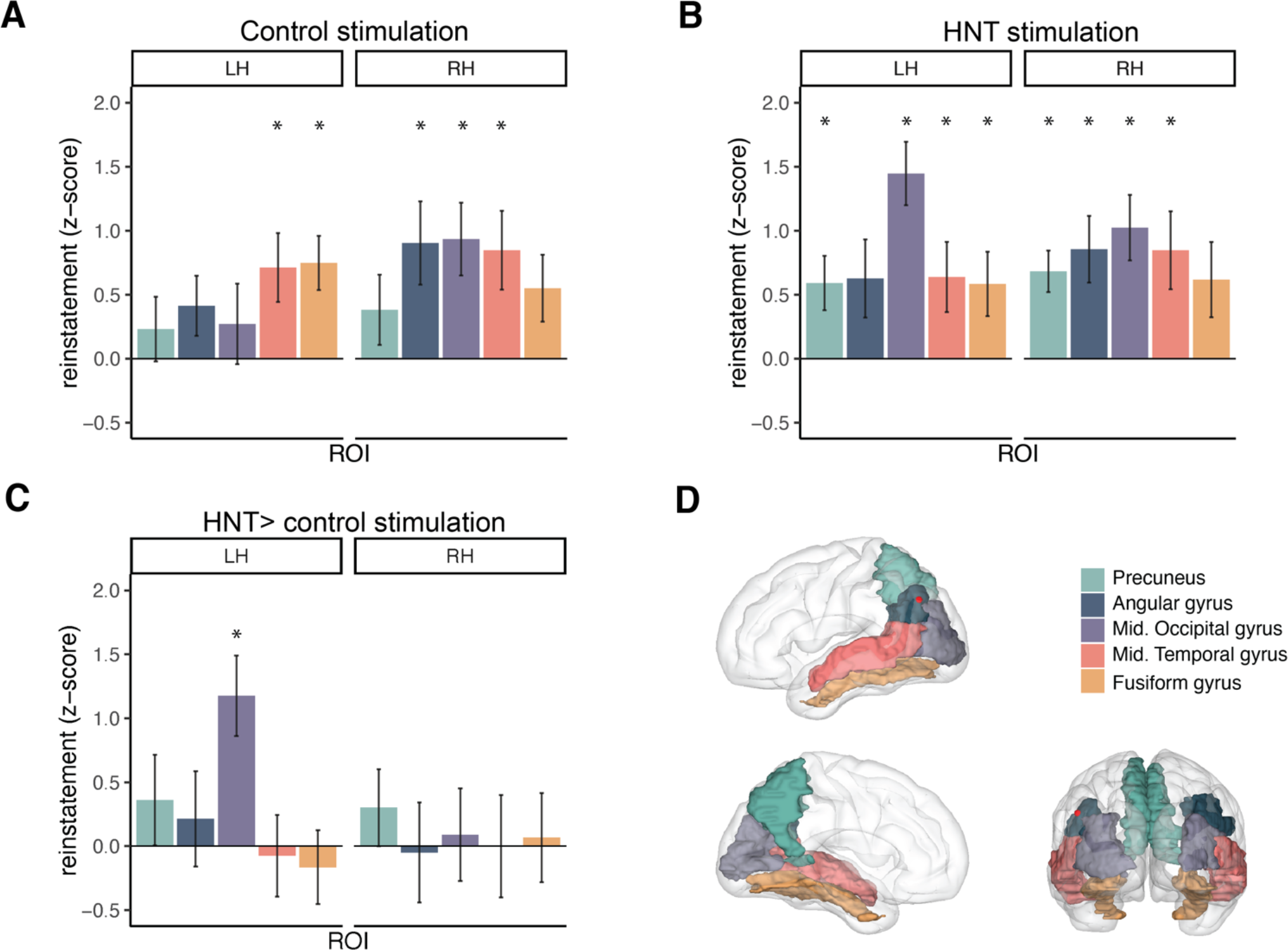
HNT stimulation increases video-specific reinstatement. (A) Reinstatement was significantly greater than zero in the indicated posterior parietal and occipital regions following control stimulation. Left (LH) and right (RH) hemispheres are plotted separately. (B) HNT stimulation led to significant reinstatement in a larger set of ROIs, including all of those showing reinstatement in control stimulation sessions. (C) Direct comparison of the two sessions (HNT – control) showing that HNT stimulation increased reinstatement in left medial occipital gyrus. Error bars represent standard error of the mean. Asterisks indicate regions that were significant after correction for multiple comparisons. (D) ROIs shown on a glass brain, with the average lateral parietal HNT stimulation location plotted as a red sphere.

For control-stimulation sessions, one-sample t-tests corrected for multiple comparisons (see Encoding-retrieval reinstatement in Methods) indicated that reinstatement was significantly greater than the null distribution for left fusiform, right middle occipital, right angular, and bilateral middle temporal ROIs (See Figure 3A; Table 2). For HNT-stimulation sessions, t-tests corrected for multiple comparisons revealed more ROIs with significant reinstatement, including all of the same ROIs with significant reinstatement in the control-stimulation condition, plus additional ROIs (left middle occipital gyrus and bilateral precuneus) (See Figure 3B; Table 2). The additional ROIs showing reinstatement following HNT stimulation relative to control stimulation were predominately in the left hemisphere, which is notable as HNT stimulation was delivered to left hemisphere based on connectivity with left hippocampus, and connectivity of posterior cortex with hippocampus is primarily unilateral [19,28].

**Table 2.** Reinstatement by ROI and stimulation condition

| Region | Mean z score (SE) |  |  | p value |  |  |
| --- | --- | --- | --- | --- | --- | --- |
|  | Control | HNT | HNT - control | Control | HNT | HNT vs. control |
| L fusiform gyrus | **0.74 (.21)** | **0.57 (.24)** | -0.17 (.29) | 0.003* | 0.032* | 0.628 |
| R middle occipital gyrus | **0.95 (.29)** | **1.04 (.25)** | 0.09 (.35) | 0.004* | <0.001* | 0.810 |
| R angular gyrus | **0.91 (.33)** | **0.88 (.27)** | -0.03 (.39) | 0.012* | 0.004* | 0.925 |
| L middle temporal gyrus | **0.70 (.27)** | **0.64 (.27)** | -0.07 (.32) | 0.017* | 0.031* | 0.754 |
| R middle temporal gyrus | **0.85 (.32)** | **0.84 (.31)** | -0.01 (.40) | 0.015* | 0.013* | 0.884 |
| L middle occipital gyrus | 0.28 (.31) | **1.45 (.24)** | **1.18 (.31)** | 0.388 | <0.001* | 0.002* |
| R precuneus | 0.38 (.27) | **0.70 (.16)** | 0.32 (.30) | 0.183 | <0.001* | 0.300 |
| L precuneus | 0.23 (.25) | **0.60 (.21)** | 0.36 (.35) | 0.362 | 0.010* | 0.318 |
| R fusiform gyrus | 0.55 (.27) | 0.63 (.29) | 0.08 (.35) | 0.054 | 0.042 | 0.828 |
| L angular gyrus | 0.41 (.23) | 0.64 (.31) | 0.23 (.37) | 0.096 | 0.054 | 0.546 |

\*reported p-value passed Benjamini-Hochberg correction for multiple comparisons
Bolded mean values are significant as measured by one-sample t-tests
Difference z-score reflects the average of within-subject differences between HNT - control

For all ROIs that showed significant reinstatement for either session, paired-samples t-tests corrected for multiple comparisons were used to test whether HNT stimulation led to greater reinstatement than control stimulation. HNT stimulation significantly enhanced reinstatement in the left middle occipital gyrus (t(19) = 3.66, p = .002, *<u>d</u>* = .936), but not in other ROIs (all p’s > 0.1). See Figure 3C; Table 2. Notably, this ROI showing greater reinstatement due to HNT stimulation is spatially distinct from the parietal location where HNT stimulation was applied (Figure 3D) and is not an element of the core hippocampal network as defined in previous experiments [13,18].

### Specificity of the effects of stimulation on reinstatement

Although the hippocampus is thought to be important for mediating cortical reinstatement [1–3], we did not predict that such reinstatement would occur within the hippocampus itself. That is, it is unclear how the dominant trisynaptic circuit within the hippocampus could exhibit patterns of activity that could support item-specific reinstatement as measured by fMRI. Further, previous findings are mixed [29], with some fMRI studies failing to find evidence for item-[4,5] or category-level [30] reinstatement within the hippocampus and other studies finding evidence for item-level [31] or category-level [32] reinstatement. We used left and right hippocampus as ROIs for the same reinstatement analysis described above, and found no evidence of significant reinstatement for either control-stimulation (p’s > 0.063) or HNT-stimulation sessions (p’s > 0.183), and no difference in reinstatement between stimulation sessions (p’s > 0.110).

To test the anatomical specificity of the effects of stimulation on reinstatement within the cortical ROIs that were used for the main analysis (Figure 3D), we performed the same analyses for ROIs that were not expected to show reinstatement effects: bilateral supplementary motor cortex and middle frontal gyrus. This ROI-based approach has similar power to the primary analyses reported above. No control regions had significant reinstatement for either the control or HNT stimulation sessions (all p’s > 0.087), and reinstatement did not differ between stimulation sessions for any region (all p’s > 0.643). These findings demonstrate that cortical reinstatement and the effects of stimulation on reinstatement were relatively specific to the ROIs selected *a priori*.

To test that the effects of stimulation on reinstatement were not merely due to different levels of univariate activity across the stimulation sessions [33], we assessed the effect of HNT stimulation on univariate activity at encoding and retrieval. We focused on the left medial occipital gyrus ROI that showed an effect of HNT relative to control stimulation on reinstatement. Voxel-wise analysis of univariate activity within this region did not differ between stimulation sessions for either encoding or retrieval (no clusters > 10 voxels at p < 0.005 threshold). We next computed the trial-wise correlation between univariate activity and reinstatement within the left medial occipital gyrus to determine whether univariate activity was driving the reinstatement effects. At encoding, average univariate activity was negatively correlated with reinstatement within this region for HNT stimulation (-t(19) = 2.98, p = 0.008), and trending towards a negative relationship for control stimulation (t(19) = −2.00, p = 0.060), but the correlation did not differ between sessions (t(19) = 1.14, p = 0.270). At retrieval, univariate activity and reinstatement were positively correlated for both HNT (t(19) = 7.48, p < 0.001), and control stimulation (t(19) = 9.43, p < 0.001), however the correlation did not differ between sessions (t(19) = −0.887, p = 0.386). Together these control analyses demonstrate that the effect of HNT stimulation on reinstatement is related to event-specific, distributed patterns of fMRI activity rather than to an overall effect on the magnitude of the fMRI response.

### Effects of stimulation on hippocampal contributions to reinstatement

To test the hypothesis that HNT stimulation would affect hippocampal activity [22], we compared univariate hippocampal activity between stimulation sessions. We focused on left hippocampus because it was the indirect target of HNT stimulation, chosen *a priori* on anatomical grounds and by indirect targeting via a lateral parietal site with high resting-state fMRI connectivity. A voxel-wise paired t-test on hippocampal activity at encoding revealed a cluster in the left posterior hippocampus showing reduced activity for HNT compared to control stimulation (7 voxels at p < 0.005, cluster corrected; peak t = −4.44) (Figure 4A). Control and HNT stimulation both showed positive activity for this cluster, although this activity was sub-threshold for HNT stimulation (control: peak t = 5.81 ; HNT: peak t = 1.29). Moreover, activity in the left hippocampus as a whole was predominately positive for both control and HNT stimulation. This suggests that HNT stimulation reduced the degree of positive hippocampal activity rather than causing negative deflections in activity. No clusters survived thresholding for the comparison between stimulation sessions on hippocampal activity at retrieval. There were no differences between stimulation sessions in the right hippocampus for either encoding or retrieval. Thus, HNT stimulation led to reduced activity at encoding, specifically in the left hippocampus which was indirectly targeted. This is consistent with previous findings of HNT stimulation’s effects on left hippocampal activity (12), and of reduced hippocampal activity due to the specific single-session theta-burst TMS parameters that were used here [25].

**Figure 4.**
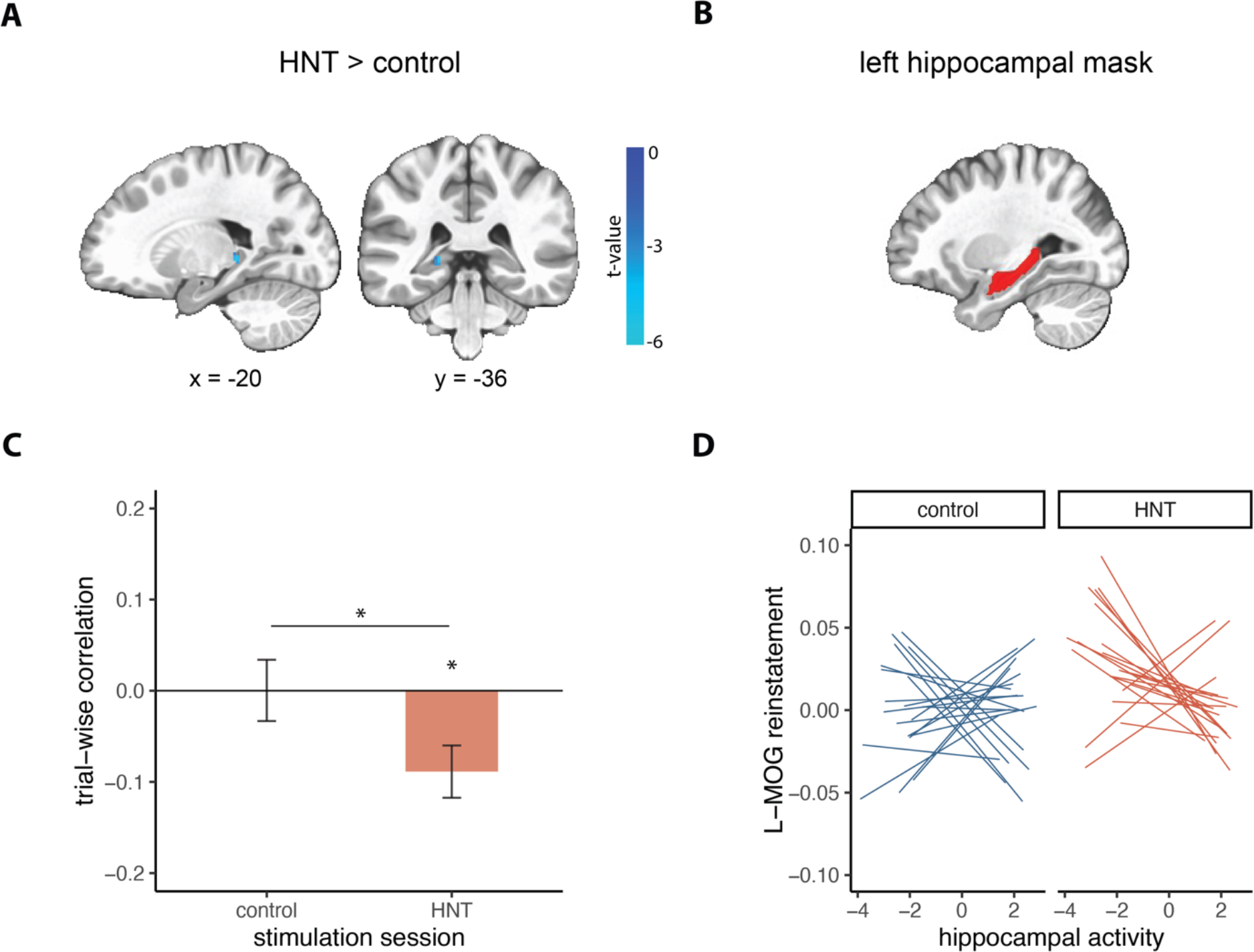
Effects of HNT stimulation on hippocampal contributions to reinstatement. (A) Univariate activity during encoding masked by the left hippocampus showed a significant cluster of negative activation for HNT vs control stimulation sessions (p < .005 cluster corrected, image thresholded at p = 0.01 for visualization purposes). Voxel-wise t-values are overlaid on a template MNI brain. (B) Left hippocampal mask used for (A), and for calculating average hippocampal activity for (C) and (D). (C) Mean within-subject (trial-wise) correlation between left hippocampal activity at encoding and reinstatement in left medial occipital gyrus (L-MOG) for control (blue) and HNT (red) stimulation sessions. Error bars represent standard error of the mean. Asterisks indicate significant comparisons (p< 0.05). (D) Within-subjects correlation between hippocampal activity and L-MOG reinstatement plotted for control and HNT stimulation sessions separately. Each line represents an individual participant’s linear fit for a given session.

To assess the relevance of HNT stimulation’s effects on hippocampal encoding activity for later reinstatement, we analyzed the trial-by-trial association between hippocampal activity and cortical reinstatement, and the effect of stimulation on this relationship. Based on the effects of HNT stimulation described above, we focused on the trial-by-trial relationship between left hippocampal univariate activity during encoding (Figure 4B) and left medial occipital gyrus encoding-retrieval reinstatement. Trial-level reinstatement was calculated for each participant by taking the difference of the Fisher transformed correlation between the diagonal value for a trial (matched videos at encoding and retrieval) and the mean of the off-diagonal correlation values for that trial (mismatched videos). Within-subject correlation coefficients were then submitted to t-tests.

For control stimulation, there was no significant relationship between univariate hippocampal activity and occipital reinstatement (t(19) = 0.02, p = 0.986, *d* = 0.004), whereas for HNT stimulation there was a significant negative association (t(19) = −3.08, p = 0.006, *d* = 0.689). 17 out of 20 participants demonstrated this negative relationship following HNT stimulation, compared to only 9 out of 20 for control stimulation. Paired-samples t-tests indicated that HNT stimulation led to a significantly more negative relationship relative to control stimulation (t(19) = 2.32, p = 0.032, *d* = 0.634). See Figure 4C and 4D. This suggests that the degree to which HNT stimulation decreased hippocampal activity during encoding was associated with increased encoding-retrieval similarity. As we will describe below, this relationship may be related to the uniqueness of the activity patterns evoked by individual videos during encoding (i.e. the degree of fMRI activity pattern overlap between similar videos at encoding).

We also examined the relationship between hippocampal activity at retrieval and cortical reinstatement [4,5]. Left hippocampal activity was positively associated with left medial occipital gyrus reinstatement for both control (t(19) = 3.95, p < 0.001, *d* = 0.883) and HNT stimulation (t(19) = 4.07, p < 0.001, *d* = 0.910). This association did not significantly differ between sessions (p = 0.784). Thus, hippocampal activity during retrieval was positively associated with reinstatement, without variation by stimulation condition, indicating that positive deflections in hippocampal activity reflect retrieval of video-specific content and that the effects of HNT stimulation were primarily on encoding.

### Testing the role of hippocampus in encoding activity that supports reinstatement

We performed an exploratory analysis to test a hypothesis about why HNT stimulation induced a negative association between left hippocampal activity during encoding and subsequent reinstatement in left middle occipital gyrus. Specifically, we reasoned that greater hippocampal activity at encoding to a specific video might partially reflect the retrieval of similar content from other videos, and that this could increase the similarity among video representations and thereby lead to reduced potential for reinstatement of specific videos. Thus, we hypothesized that greater hippocampal activity at encoding would be related to higher similarity of fMRI activity patterns between videos at encoding, and that this “encoding similarity” would in turn be related to reduced video-specific cortical reinstatement at retrieval. Encoding similarity was calculated as the average Fisher-transformed correlation between the pattern of fMRI activity for each video and all other videos at encoding (See Encoding similarity in Methods).

Within the left medial occipital gyrus, encoding similarity was significantly greater than zero for control (t(19) = 14.16, p < 0.001) and HNT sessions (t(19) = 14.49, p < 0.001) and did not differ by stimulation session (t(19) = 1.68, p = 0.109). Although they were not the focus of this exploratory analysis, we also assessed encoding similarity within each of the 5 bilateral ROIs (Figure 3D). Values for all ROIs were significantly greater than zero for both stimulation sessions (all p’s < 0.001) with no significant differences between stimulation sessions (all p’s > 0.05, corrected for multiple comparisons), however all ROIs showed a trend towards lower encoding similarity for HNT versus control stimulation. This high similarity of activity patterns across videos is unsurprising and likely reflects similarity in video content and task demands across trials. The critical question of interest is thus whether encoding similarity is related to univariate hippocampal encoding activity and subsequent pattern reinstatement.

Indeed, there was a significant within-subject correlation between encoding similarity and hippocampal activity at encoding, with greater hippocampal activity associated with higher encoding similarity in left medial occipital gyrus for both control (t(19) = 9.42, p < 0.001) and HNT sessions (t(19 = 5.37, p < 0.001), with no difference between sessions (t(19)= −0.40, p = 0.692) (Supplementary Figure 1A). Thus, greater hippocampal activity was associated with more overlap of fMRI activity patterns among different videos in occipital cortex at encoding.

As hypothesized, higher encoding similarity was in turn related to less cortical reinstatement. There was a significant negative within-subjects correlation between encoding similarity and reinstatement for both control (t(19) = −3.98, p < 0.001) and HNT stimulation sessions (t(19)= −3.69, p = 0.002), with no difference by stimulation session (t(19 = 0.64, p = 0.532) (Supplementary Figure 1B). Together these findings suggest that lower hippocampal activity was generally related to more dissimilarity among video activity patterns during encoding, and that more dissimilar encoding patterns in turn predicted greater video-specific activity reinstatement during retrieval. The beneficial effect of HNT stimulation on reinstatement thus could have resulted from its suppressive effect on hippocampal activity during encoding.

## Discussion

This study tested the effect of HNT stimulation on memory for naturalistic episodes. We found that stimulation of parietal regions showing high functional connectivity with the hippocampus improved memory accuracy and induced reinstatement in a greater number of posterior cortical regions, significantly enhancing reinstatement in the middle occipital gyrus. We further found that HNT stimulation led to reduced univariate activity in the left hippocampus at encoding, and identified a stimulation-induced negative relationship between trial-wise hippocampal activity and cortical reinstatement that appeared to be mediated by the neural distinctness of videos at encoding. These findings demonstrate that perturbing one node of the episodic memory network can alter large-scale, multimodal patterns of neural activity which are considered to be fundamental to memory.

Episodic memory is thought to critically depend on cortical reinstatement mediated by the hippocampus [1–3]. Consistent with prior research [9,27,34], we identified reinstatement of video-specific patterns of neural activity within a set of posterior parietal and occipital regions following control stimulation. A greater number of regions showed significant reinstatement following HNT stimulation, and many of these regions were located in the left hemisphere, suggesting that left lateralized stimulation preferentially affects cortical areas in the left hemisphere. Direct comparison of the two sessions revealed that HNT stimulation led to greater reinstatement in the left middle occipital gyrus, a region implicated in episodic memory encoding success [35], recall of visual images [36], and reinstatement of video clips [9,27]. This region might be effected by stimulation due to its proximity to the left lateral HNT stimulation site (Figure 3D) and its reported functional connectivity with the hippocampus during memory formation [15] and retrieval [37], Thus, we speculate that the medial occipital gyrus may be particularly responsive to stimulation in the present study due to its established role in neural reinstatement, episodic memory encoding, and visual memory retrieval, as well as its connections with the hippocampus. Importantly, these findings demonstrate that HNT stimulation alters reinstatement in posterior parietal and occipital regions typically not considered to be part of the core hippocampal network.

We found that a single session of theta-burst HNT stimulation improves memory accuracy and enhances cortical reinstatement of naturalistic episodic memories. These findings add to prior research which identified improvements in classically hippocampal-dependent word-list [23] and spatial precision tests [24] of episodic memory using the same stimulation paradigm, with associated increases in resting-state fMRI connectivity between the hippocampus and regions of its core network [23]. Our findings of enhanced episodic memory accuracy and reinstatement are consistent with the improvements identified in these [23,24] as well as other HNT stimulation studies using a multi-day paradigm [38–41]. Conversely, theta-burst HNT was recently shown to reduce evoked hippocampal activity and performance on a task thought to rely on episodic memory [25]. Others have similarly reported disrupted episodic memory following stimulation of lateral parietal regions [25,26,42–46], although these studies did not use an HNT stimulation approach. Notably, the studies showing disruptions tested the effects of stimulation on memory processes thought to be partially dependent on the parietal lobe, such as multimodal integration [42], generation [25] and recall of detailed multisensory information [26,43], and memory vividness [44], which differ from the predominately hippocampal-dependent measure of memory in ours and similar studies [24,38–41,47]. This pattern of results is consistent with a recent proposal that stimulation may disrupt local processing at the site of stimulation, while enhancing processing in downstream, functionally connected regions such as the hippocampus [22]. Thus, discrepancies in disruptive versus faciliatory effects of stimulation on memory may be due to differences in the neural basis of the targeted memory function.

Prior work has shown that delivering TMS to accessible portions of the episodic memory network can alter performance on traditional laboratory tests of episodic memory tasks [23,24,38,41,48]. The present findings demonstrate that this approach can be used to improve objective memory for complex stimuli that more closely approximate memory demands experienced outside the laboratory. We found that HNT stimulation altered objective memory accuracy on the second of two true/false questions. Interestingly, participants performed worse on the second question than they did for the first question, but only for control stimulation. One possible explanation for this finding is that HNT stimulation leads to stronger reactivation of a given video during the mental imagery phase of a trial, making the memory more temporarily stable, which allows participants access to the details of a memory for longer, allowing them to answer the second question more accurately. While the present behavioral data is not sufficient to formally test this idea, future studies could experimentally test whether HNT stimulation affects the temporal stability of a reactivated memory.

We stimulated areas of the lateral parietal lobe with high functional connectivity to the hippocampus, aiming to indirectly target the hippocampus, and found that HNT stimulation led to reduced univariate activity at encoding in a small left posterior hippocampal cluster. At the trial-level, HNT stimulation led to a significant negative association between left hippocampal activity at encoding and reinstatement in the left medial occipital gyrus. Follow-up analyses revealed that this negative association was related to the similarity or interference between videos as they were being encoded. Greater neural similarity in the left medial occipital gyrus between different videos at encoding was associated with concurrent increases in hippocampal activity and lower cortical reinstatement during retrieval. This suggests that hippocampal activity during encoding might partially reflect retrieval of content from other videos, which in turn makes later reinstatement of the unique elements of an individual video more difficult. The stimulation-induced reduction in hippocampal activity at encoding might therefore reflect less neural overlap (and interference) between similar videos at encoding, which in turn leads to greater reinstatement of unique videos. Indeed, we found a trend towards lower encoding similarity following HNT stimulation in all of the ROIs measured, although these effects were nonsignificant. Future studies are needed to directly test the effect of stimulation on the relationship between hippocampal activity, encoding similarity, and neural reinstatement.

To conclude, the present findings demonstrate that noninvasive stimulation of functionally connected regions can alter large-scale patterns of event-specific activity beyond the site of stimulation, leading to enhanced memory for complex, naturalistic events and their spatiotemporal contexts. Together with prior findings of altered episodic memory and neural activity following network-targeted stimulation [23,24,38,41,48], our findings support the ability of HNT to alter hippocampal-cortical networks important for memory.

## Materials and Methods

### Participants

Twenty adults participated in the current study (11 females, mean age = 24.26, SD = 3.57, range = 19-32). All participants were native or fluent English speakers, had normal or corrected-to-normal vision, and were free from a history of neurological illness or injury, psychiatric condition, substance abuse, or serious medical conditions. All participants passed standard MRI and TMS safety screenings [49]. Participants provided informed consent prior to participating in the experiment and were paid for participation. Study procedures were approved by the Northwestern University Institutional Review Board.

### Experimental design

Participants completed a baseline session and two experimental sessions on separate days in a within-subjects design. During the baseline session, resting-state fMRI and structural MRI were acquired, and TMS resting motor threshold was determined. Resting-state fMRI from this session was used to define the stimulation locations for subsequent sessions. Each experimental session lasted approximately 2 hours and occurred on separate days, at least 48 hours apart (mean = 3.1 days, range = 2-7). In experimental sessions, participants received continuous theta burst TMS to either the control site (vertex) or the lateral parietal target (HNT), with the order of stimulation site counterbalanced across participants. TMS was administered in a room adjacent to the MRI scanner room. Immediately following the final TMS pulse, participants were moved to the scanner where they performed an episodic memory task while fMRI was collected. See Figure 1B for an overview of experimental sessions. During the encoding phase of the episodic memory task, participants watched short video clips, and at retrieval they recalled the videos, answered true/false questions to test their accuracy for the videos, and rated their subjective vividness of the memory. The encoding phase of the memory task began on average 6.9 mins (SD = 1.1, range = 5-10 mins) after the end of TMS delivery and was completed within approximately 21 minutes (length of encoding phase = 14 mins) post-TMS. Retrieval began immediately after encoding and was completed within approximately 55 mins of the final TMS pulse (length of retrieval phase = 32 mins).

### Subject-specific stimulation target identification

HNT lateral parietal stimulation locations were identified separately for each participant based on resting-state fMRI connectivity with a left hippocampal seed, measured during the baseline session. This procedure was adapted from previous studies from our laboratory [41,48]. Baseline resting-state and structural MRI data were preprocessed using AFNI [50]. Preprocessing steps included motion correction, slice-timing correction to the first slice, functional/structural co-registration, resampling to a resolution of 1.5×1.5×1.5 mm, stereotactic transformation using Montreal Neurologic Institute 305 (MNI-305) template, band-pass filtering (0.01-0.10Hz), spatial smoothing (with a 4-mm FWHM Guassian kernel), despiking, linear detrending, and regressing out the motion timeseries. Participant-specific hippocampal coordinates were located by identifying a voxel in the left hippocampal body nearest to MNI [-29,-25,-13] that demonstrated high connectivity with bilateral hippocampus. This hippocampal voxel was then used as a seed in a whole-brain resting-state functional connectivity analysis. We then identified clusters of voxels in left parietal cortex with high functional connectivity to the hippocampal seed that fell within left angular and supramarginal gyri, nearest to MNI [−47, −68, 36]. Left parietal TMS target coordinates were chosen as the peak of the parietal cluster that fell within ~2cm of the skull and was located on a gyrus (mean correlation between seed-target = .38, range = .23-.48). As a control site, the vertex was located on each participant at the MNI coordinate [0,-15,74], and adjusted slightly so it fell within the interhemispheric fissure. HNT and control stimulation locations were transformed from MNI space into each participant’s native MRI space for anatomically guided TMS. Individual hippocampal seeds, lateral parietal targets, and vertex targets are plotted in Figure 1A.

### TMS procedure

Participants received TMS to the left parietal target or to the control vertex site on separate experimental days. Resting motor threshold (RMT), determined during the baseline session, was defined as the minimum stimulator output necessary to produce a visible contraction of the right thumb (abductor pollicis brevis) for 5 out of 10 consecutive single biphasic pulses delivered to the left M1 thumb area. Continuous theta burst stimulation was applied at 80% RMT for both stimulation locations. The Localite (GmBH, Germany) frameless stereotactic system was used to target the selected stimulation locations. Five anatomical landmarks located on the face were used to co-register the anatomical MRI to the participant’s head, and landmarks on the scalp were used to improve registration based on the curvature of the participant’s head. An infrared camera recorded sensors attached to the participant and the TMS coil, allowing for real-time tracking of the TMS coil over the participant’s MRI. A MagPro X100 stimulator connected to a Cool-B65 butterfly coil (MagVenture A/S, Farum, Denmark) was used to deliver stimulation. The continuous theta burst stimulation protocol consisted of a total of 600 biphasic pulses arranged in bursts delivered every 200 ms (5-Hz), with each burst containing three pulses delivered at 50Hz (40 s duration). The coil was positioned perpendicular to the brain surface at the stimulation site with the magnetic field oriented across the gyrus.

### Episodic memory task

Practice sessions were performed prior to TMS administration to familiarize participants with the task and to ensure correct performance. Once moved to the MRI scanner, participants completed the episodic memory task in which they watched and recalled a series of short video clips. Videos consisted of 102 short (7 s) depictions of common events such as draining pasta, kicking a ball, and putting a sheet on the bed. Each video depicted a unique event centered around the same one character and occurred in a fixed number of locations, resulting in high overlap between elements of the videos *(*note videos will be made available in an archive)*. All videos were presented without sound. Videos were divided into 2 lists of 51 for presentation on separate experimental sessions, the order of which was counterbalanced across days and stimulation conditions. Within each list, videos were presented in a randomized order at both encoding and retrieval phases.

At the start of the encoding phase, participants were reminded of the task instructions and were instructed to pay close attention to all elements of the videos as their memory for the videos would later be tested. Each video was then presented alone for 7 s, followed by an interstimulus interval (ISI) of 1 s. Participants then judged whether a number was odd or even for 2 s to discourage continued processing of the video. Previous evidence suggests that such judgments drive activity in key regions such as hippocampus back to zero during the intertrial interval (ITI) [51]. The trial ended with an ITI (fixation cross), jittered at 4, 6, and 8 s. Each encoding trial was 16 s on average. All encoding trials occurred within 1 fMRI run lasting 13.9 mins.

The retrieval phase began immediately following completion of the encoding phase. Following a reminder of the instructions for this phase, participants were presented with the description of a video for 3 s and told to mentally replay the video within the allotted time, within a blank box that remained on the screen alone for 7 s. After mentally recalling each video, participants rated their vividness of the memory on a scale from 1-4, with 1 meaning they did not recall the video at all, and 4 meaning they recalled it vividly. 4 s were allotted for vividness ratings. Following the vividness rating, participants answered two true/false (yes/no) accuracy questions about the videos, incorporating their confidence (1-definitely yes, 2-maybe yes, 3-maybe no, 4-definitely no). 5 s were allotted for each accuracy question. There was a total of 1.2 s ISI within each trial, and an ITI (fixation cross) jittered at 4, 6, and 8 s, with an average trial length of 34 s. Responses were made using 2 response boxes, each with 2 buttons. Retrieval trials occurred over 3 fMRI runs (17 trials each), each lasting 10 mins.

### MRI acquisition

Structural and functional images were acquired using a Siemens 3 T Prisma whole-body scanner with a 64-channel head coil located in the Northwestern University Center for Translational Imaging Facility. Baseline resting-state and functional images during experimental sessions were acquired using a whole-brain BOLD EPI sequence (TR = 2000 ms, TE = 20 ms, FOV = 1116×1080 mm, flip angle = 80°, and 1.7×1.7×1.7 mm voxel resolution, over 275 volumes). Structural images were acquired using a T1-weighted MPRAGE sequence (TR = 2170 ms, TE = 1.69 ms, FOV = 256×256 mm, flip angle = 7°, voxel resolution: 1.0 × 1.0 × 1.0 mm, 1-mm thick sagittal slices). During the baseline resting-state scan, participants were instructed to lie still with their eyes open. The baseline resting-state scan was 275 volumes (9.2 mins). The encoding phase of the memory task consisted of one 13.9 min run (418 volumes), while retrieval was split among three 10 min runs (299 volumes each).

### Data analyses

#### Behavioral analyses

To determine the effect of HNT stimulation on objective episodic memory, we conducted a 2×2 repeated-measures ANOVA with accuracy as the dependent variable, and question number (Q1 vs Q2) and stimulation session (HNT vs control) as factors. Follow-up paired-samples t-tests were performed to assess significant interactions and main effects. Effect sizes were reported for primary behavioral outcomes. Confidence ratings were extracted from accuracy ratings, such that responses 1 and 4 reflected high confidence and responses 2 and 3 reflected low confidence. Paired-samples t-tests were conducted comparing HNT to control stimulation on vividness and confidence ratings. Trials were excluded from behavioral analyses if there was no response to either of the three retrieval questions. On average, 97% of trials were included for HNT stimulation sessions, and 98% of trials were included for control stimulation sessions.

#### MRI preprocessing

Functional MRI data for experimental sessions were preprocessed using AFNI software. Preprocessing included functional-structural co-registration, motion correction, spatial smoothing using a 1.7-mm full-width-half-maximum (FWHM) isotropic Gaussian kernel, and signal intensity normalization by the mean of each voxel. Motion parameters were calculated for each volume, and volumes with excessive frame to frame displacement (>0.3 mm) were flagged for later censoring.

Single-trial estimates were generated for multivariate analyses using a general linear model (GLM) in AFNI (*3dDeconvolve*). A separate model was constructed for each individual trial to estimate its’ activity separately from all other trials and nuisance variables, an approach known to work effectively for single-trial estimation for multivoxel pattern analyses [52]. For each functional run, individual trials were modelled separately against all other trials using a response model of a 7-s block convolved with a canonical hemodynamic response function. Nuisance variables included the six affine motion estimates generated by motion correction as well as linear drift. For encoding trials, the 7-s block began at the start of video presentation, while at retrieval the 7-s block began when participants saw the cued video title. The resulting single-trial t-maps for each trial were used for subsequent analyses, based on recommendations that using t-maps rather than beta maps for representational similarity analyses reduces the influence of noisy voxels [33]. All multivariate analyses were carried out in native space.

For univariate analyses, encoding- and retrieval-related neural activity were separately modelled for all trials using 7 s block functions convolved with a hemodynamic response function in addition to six motion regressors. Neural activity associated with accurate trials (measured by accuracy on the second question) were modelled separately from inaccurate trials, and then added together to represent all trials. For both sets of GLMs, trials were excluded from analyses if there was no response to the odd/even question at encoding, or no response to either of the three retrieval questions. This resulted in inclusion of an average of 97% trials for control sessions, and 96% trials for HNT sessions.

#### Regions of interest

Multivariate analyses were focused on a set of ROIs. ROIs included bilateral precuneus, angular gyrus, middle occipital gyrus, middle temporal gyrus, and fusiform gyrus. These regions were chosen because they are known to show reinstatement effects of video-specific patterns of activity [9]. ROIs were defined using the Eickhoff-Zilles macro labels from N27 in MNI space (CA_ML_18_MNI atlas in AFNI) on a template brain in MNI space and warped to native space for each participant by calculating the transformation matrix needed to warp into MNI space and then reverse-transforming all ROIs into native space using affine transformation (*3dAllineate*).

#### Encoding-retrieval reinstatement

We examined reinstatement of video-specific patterns of neural activity by conducting a series of representational similarity analyses (RSAs) on patterns of neural activity within ROIs. RSAs measured encoding-retrieval similarity by computing the correlation between neural activity patterns for all pairs of videos, for each ROI. Only trials that were self-reported as recollected (vividness score of > 1) were included in these analyses (86% trials for control; 85% trials for HNT). Pairwise correlations between all videos resulted in a matrix with the diagonal reflecting correlations between the same video at encoding and retrieval (matched pairs), and the off-diagonal reflecting correlations between different videos (mismatched pairs). Pairwise correlation values were Fisher transformed. We then subtracted the mean of the off-diagonal correlation values from the mean of the diagonal correlations, resulting in a metric that reflects the degree to which neural pattern similarity was greater for matched versus mismatched video pairs. This metric was then z-scored against a null distribution of correlation difference scores, calculated by randomly shuffling trials over 1000 permutations. Group-level statistics were performed on these z-scored pattern similarity values. RSAs were computed in MATLAB (The MathWorks, Inc., Natick, MA, USA), using the CoSMoMVPA toolbox (cosmomvpa.org)[53] functions to load and organize fMRI datasets (cosmo_fmri_dataset), and calculate encodingretrieval similarity matrices (cosmo_correlation_measure). All subsequent analyses on correlation matrices were performed using in-house MATLAB scripts.

To assess the presence of neural reinstatement of video-specific patterns of neural activity, we first ran one-sample t-tests on z-scored pattern similarity values for each ROI, within each session. We next ran a series of paired-samples t-tests to determine the effect of stimulation on reinstatement. Only ROIs that showed reinstatement significantly greater than zero for individual sessions were entered into paired-samples t-tests. All p-values were submitted to Benjamini-Hochberg correction for multiple comparisons [54] using a q of .05, and only p-values that passed this correction are listed as significant. Effect sizes were reported for primary outcomes.

#### Univariate analyses

To determine whether HNT stimulation alters hippocampal activity, maps for both stimulation sessions were submitted to paired t-tests. Maps were first warped from native space to MNI space using affine transformation (*3dAllineate*), and masked by the left hippocampus, defined using the CA_ML_18_MNI atlas. The voxel-wise threshold was p < .005, with cluster size determined based on the spatial smoothness of the data using Monte Carlo simulations from the 3dClustSim tool in AFNI using the spatial autocorrelation function (ACF). Based on this procedure, the minimum cluster size at p = .005 for a combined threshold of p < 0.05 was determined to be 5 voxels. All univariate analyses were performed in AFNI.

#### Association between hippocampal activity and cortical reinstatement

We next examined the association between left univariate hippocampal activity and reinstatement in the left medial occipital gyrus. Average hippocampal activity was calculated for individual subjects by taking the average across all voxels encompassing the left hippocampus (CA_ML_18_MNI ROI reverse-transformed into native space using affine transformation). For each participant, we computed the trial-level correlation between mean hippocampal activity and reinstatement of a given trial, defined as the correlation difference between the diagonal value for a trial (matched videos at encoding and retrieval) and the mean of the off-diagonal values for that trial (mismatched videos). Correlation values were submitted to one-sample t-tests within stimulation sessions to determine if the association between univariate activity and reinstatement is greater than zero, and paired-sample t-tests were used to assess differences between stimulation sessions.

#### Encoding similarity

We performed a series of exploratory analyses examining similarity between patterns of neural activity for different videos at encoding. Encoding similarity was measured by computing the dissimilarity matrix between each video and all other videos at encoding using the cosmo_dissimilarity_matrix function in CoSMoMVPA [53]. This function computes the correlation distance between patterns of neural activity, resulting in a correlation matrix reflecting the dissimilarity of each video to every other video. Dissimilarity values were subtracted from 1 to reflect similarity, or the correlation between videos, and Fisher transformed. Each video/trial’s average correlation with all other trials was calculated across a row, including values in the upper triangle of the correlation matrix only, and excluding the diagonal value. We computed the within-subjects correlation between encoding similarity values and neural reinstatement in the left medial occipital gyrus, measured as the correlation difference between matched videos and the mean of mismatched videos for that trial. Finally, we computed the within-subjects correlation between encoding similarity and univariate left hippocampal activity. All correlations were performed using encoding similarity within the left medial occipital gyrus. Only trials that were self-reported as recollected (vividness score of > 1) were included in these analyses.

## Acknowledgments

We thank Stephanie Wert, Brennan Durr, and Erica Karp for their assistance with data collection. This research was supported in part through the computational resources and staff contributions for Quest, the high-performance computing facility at Northwestern University, which is jointly supported by the Office of the Provost, the Office for Research, and Northwestern University Information Technology. This research was supported by R01-MH106512 from the National Institute of Mental Health. The content is solely the responsibility of the authors and does not necessarily represent the official view of the National Institutes of Health.

## Author Contributions

M.H., J.L.V., and T.K. designed research; M.H. performed research; M.H. and J.L.V. analyzed data, M.H., J.L.V, T.K., and J.E.K. wrote the paper.

## Supplementary material

**Figure S1.**
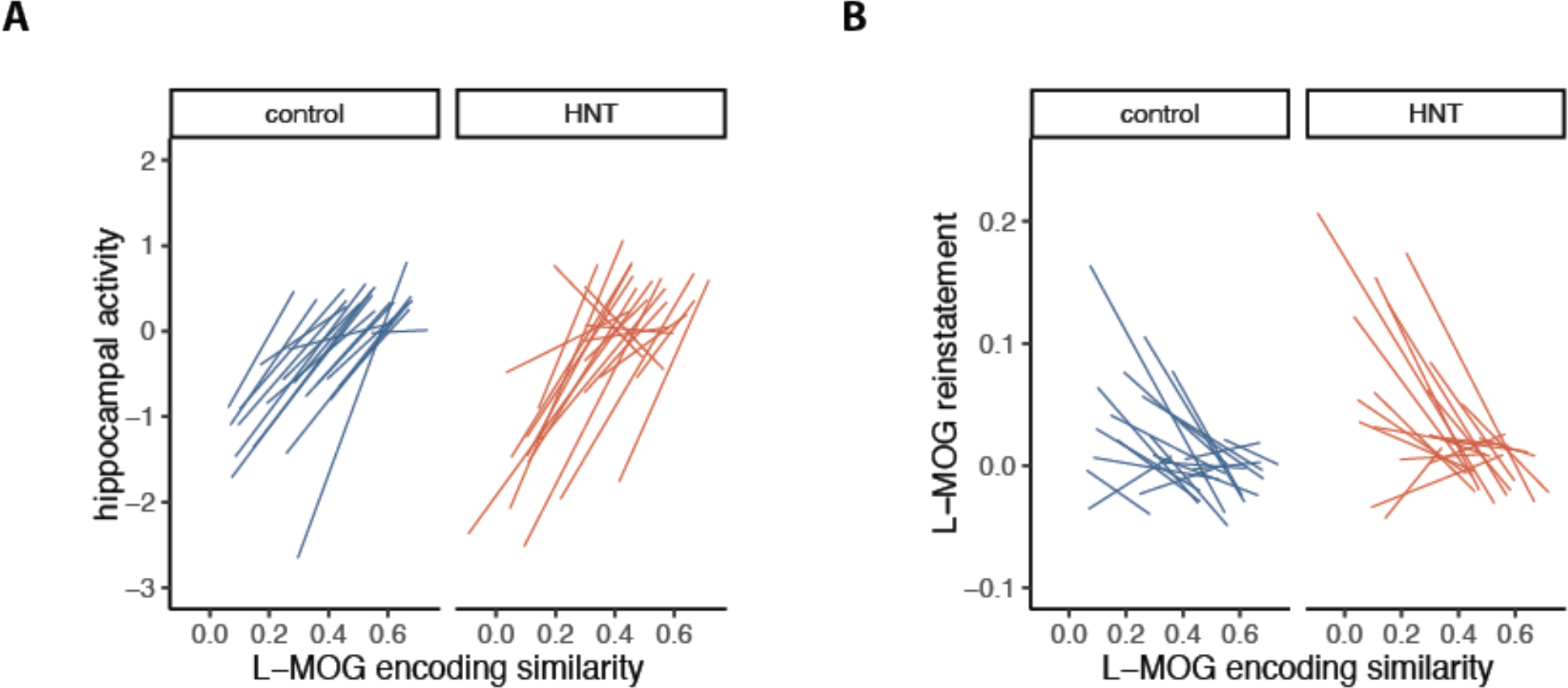
Effects of encoding similarity on hippocampal contributions to reinstatement. (A) Within-subjects correlation between average left hippocampal activity at encoding and encoding similarity in left medial occipital gyrus for control and HNT stimulation plotted separately. (B) Within-subjects correlation between encoding similarity and reinstatement in left medial occipital gyrus for control and HNT stimulation. Each line represents an individual participant’s correlation for a given session.

